# SBOL Visual 2 Ontology

**DOI:** 10.1101/2020.01.24.918417

**Authors:** Göksel Misirli, Jacob Beal, Thomas E. Gorochowski, Guy-Bart Stan, Anil Wipat, Chris Myers

## Abstract

Standardising the visual representation of genetic parts and circuits is vital for unambiguously creating and interpreting genetic designs. To this end, an increasing number of tools are adopting well-defined glyphs from the Synthetic Biology Open Language (SBOL) Visual standard to represent various genetic parts and their relationships. However, the implementation and maintenance of the relationships between biological elements or concepts and their associated glyphs has to now been left up to tool developers. We address this need with the SBOL Visual 2 Ontology, a machine-accessible resource that provides rules for mapping from genetic parts, molecules, and interactions between them, to agreed SBOL Visual glyphs. This resource, together with a web service, can be used as a library to simplify the development of visualization tools, as a stand-alone resource to computationally search for suitable glyphs, and to help facilitate integration with existing biological ontologies and standards in synthetic biology.

**Graphical TOC Entry:** 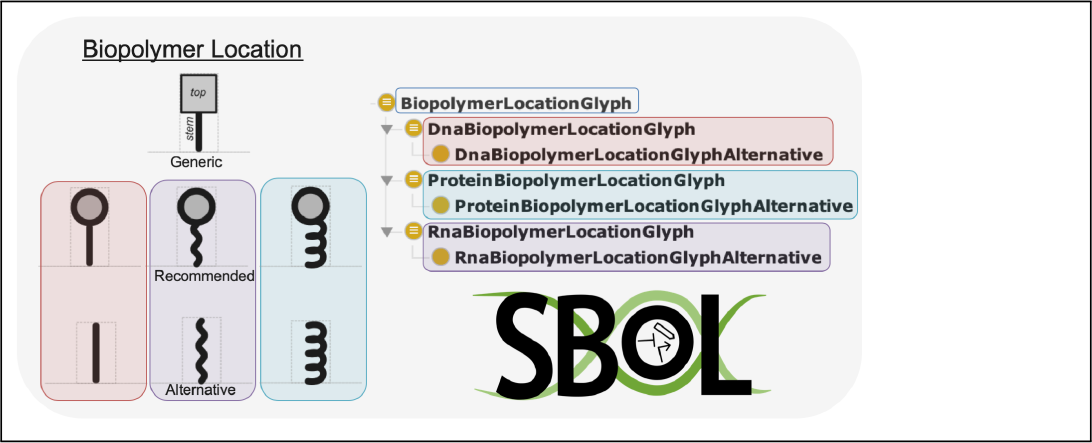

## Introduction

Visual representation of genetic designs can aid human comprehension and improve the dissemination of complex information. As genetic designs grow ever more complex, there is also an increasing need for computational tools to visualize them. The Synthetic Biology Open Language (SBOL) Visual standard^1–4^ has been developed to provide guidelines on how features of a genetic design should map to suitable glyphs, as well as how various glyphs can be connected together to unambiguously convey forms of interaction. SBOL Visual is being used by an increasing number of computational tools and repositories for synthetic biology.^3^ To date, however, the actual computational implementation of mapping from genetic parts, molecules, and interactions to suitable glyphs has been left entirely up to the tool developers. This is a significant burden, opening up the opportunity for errors to arise and causing an ongoing maintenance cost as the set of glyphs grows and evolves.

A common solution to problems of this sort is to create an ontology. An ontology is a formal representation of entities in a domain or subject. Entities are typically represented as a set of concepts (expressed as terms) that shows the entity properties and the relations between them. Ontological terms are a powerful way to represent a large amount of machine-readable information via simple URIs, which can then further point to additional properties of ontological terms and the relationships of such terms with other biological concepts. As a result, an ontological representation of SBOL Visual is highly desirable for computational processing of visual guidelines and further integration with other ontologies and tools. Indeed, an ontology was previously created for SBOL Visual 1,^5^ providing mappings between glyphs representing DNA features and terms in the Sequence Ontology.^6^ Likewise, an ontology called SBOL-OWL^7^ was recently released, providing semantic meaning for terms in the SBOL 2 data standard^8,9^ in a machine-readable format. Compared to the previous version, SBOL Visual 2 has greatly expanded in scope, matching the SBOL 2 data standard with new classes of glyphs and relationships between them, as well as many more glyphs for biological features. As a result, the SBOL Visual Ontology required an overhaul to keep pace with these developments.

To address this, we developed the SBOL Visual 2 Ontology, which provides a machine-readable representation of the constraints attached to genetic circuit glyphs and their relationships to other ontological terms. Using an ontological mapping, the SBOL Visual Ontology further integrates information about standardized glyphs with SBOL-OWL and the SBOL data standard. This information can be leveraged to simplify the implementation and maintenance of any genetic design visualization tool, as we demonstrate with an ontology-based image web service.

## Results and Discussion

The SBOL Visual 2 Ontology (hereafter abbreviated SBOL-VO) is a machine-readable representation of the SBOL Visual specification that facilitates computational access to standardised glyphs. The ontology consists of terms for different glyph types. These terms are represented as OWL classes and can include subclasses for recommended and alternative glyphs. The ontology also captures how these terms are related to each other and terms from other ontologies that are utilised by the SBOL Visual standard.

The basis for this ontology is the distribution of the SBOL Visual specification maintained by the SBOL community. The bulk of this specification comprises a set of standard glyphs for a variety of commonly used genetic parts, organized as “families” of recommended glyphs plus generic and alternative glyphs where necessary. For each glyph family, the specification is presented as a directory of glyph files and a single human-editable Markdown file in a standard format. This Markdown file includes information about the mapping between glyphs and relevant genetic design components through biological roles and molecular interactions. These mappings are identified via commonly used ontological terms in the Sequence Ontology (SO)^6^ and the Systems Biology Ontology (SBO).^10^

To capture this information, the base class in the ontology is `Glyph`, subclasses of which correspond to either an individual glyph or a family of related glyphs (Figure 1). Each Glyph class is decorated with machine-readable properties, while glyph-specific properties are captured as annotations. Specifically, a class representing a glyph may include the following `Annotation` properties: `rdfs:label` (name), `rdfs:comment` (description), `defaultGlyph` (the name of the glyph file), `glyphDirectory` (the folder containing the glyph), `notes` (additional free text information), `recommended` (whether the glyph is recommended or not), and `prototypicalExample` (an example use of the glyph).

**Figure 1:**
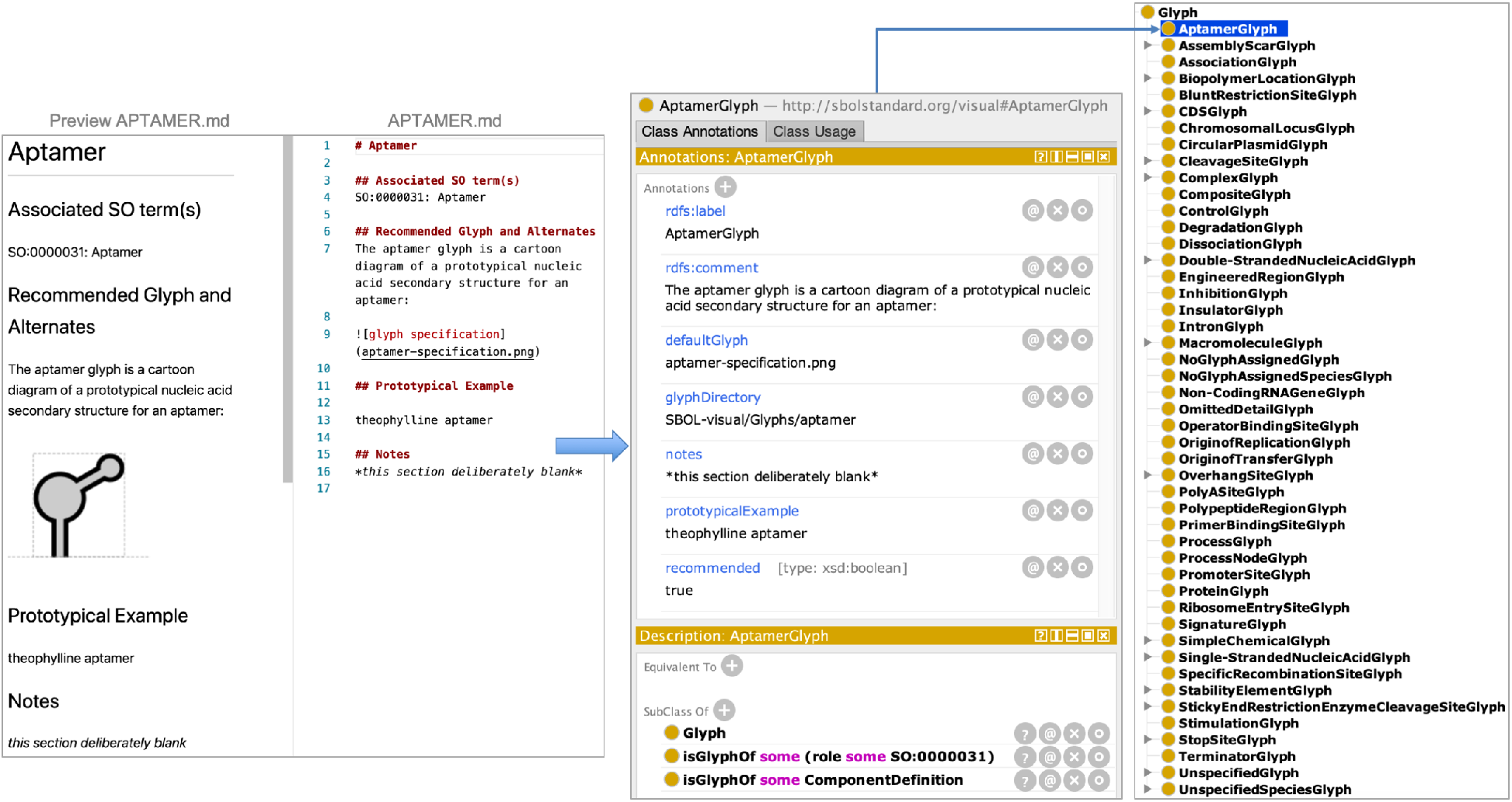
Standard-format Markdown files in the SBOL Visual distribution that describe recommended and alternative glyphs are used to create ontology classes (or terms) and restrictions.

SBOL-VO classifies glyphs into three categories to represent nucleic acid sequence features, molecular interactions, and different types of biological molecule participating in these interactions. The SBOL Visual specification already associates glyphs with terms from external ontologies such as SO and the SBO to map glyphs to biological roles. However, this information is only available as free-text. SBOL-VO makes these relations machine-readable and integrates data from external ontologies using restrictions based on the ontological terms associated with a glyph.

SBOL-VO is also integrated with the SBOL 2 data standard, which defines a format for exchange of genetic design information. These restrictions are defined for SBOL entities that represent genetic parts or molecular interactions via `sbol:ComponentDefinition` or `sbol:Interaction` respectively. For example, `AptamerGlyph` is defined to be a glyph for `sbol:ComponentDefinition` entities with the `sbol:role` of `SO:0000031`, which is an SO term used for aptamers. The following rules are applied to create mappings between SBOL-VO and other ontologies:

- If a glyph is associated with an SO term, a restriction is created to map the glyph with sequence feature entities. This restriction links the corresponding SBOL-VO class to `ComponentDefinition` entities using the role property. For example, AptamerGlyph `isGlyphOf some ComponentDefinition` with the `role of SO:0000031`.
- If a glyph is associated with an SBO term that is describing a material entity (i.e., a child term of SBO:0000240), a restriction is created to map the glyph with molecule types, such as protein or complex. This restriction links the corresponding SBOL-VO class to `ComponentDefinition` entities using the `type` property. For example, ComplexGlyph `isGlyphOf some ComponentDefinition` with a type of `SBO:0000253`.
- If a glyph is associated with an SBO term that is describing a biological activity or a process (i.e., a child term of `SBO:0000231`), a restriction is created to map the glyph with interaction entities. This restriction links the corresponding SBOL-VO class to `Interaction` entities using the `type` property. For example, DegradationGlyph `isGlyphOf some Interaction` with the type of `SBO:0000179`.

In addition, SBOL-VO defines terms to identify recommended and alternative glyphs, and the relationships between these. These terms are derived from the Markdown files, each of which is maintained for a single SBOL Visual category (e.g., ‘Aptamer’ or ‘Biopolymer Location’) and is available as free-text (Figure 1). Based on the representation of information in these Markdown files, hierarchical relationships between SBOL-VO terms are created. For each SBOL Visual category, there may be one or more glyphs defined. When the mapping between a category and a glyph is one-to-one, a single glyph term is defined representing the category (e.g., AptamerGlyph, see Figure 2A).

**Figure 2:**
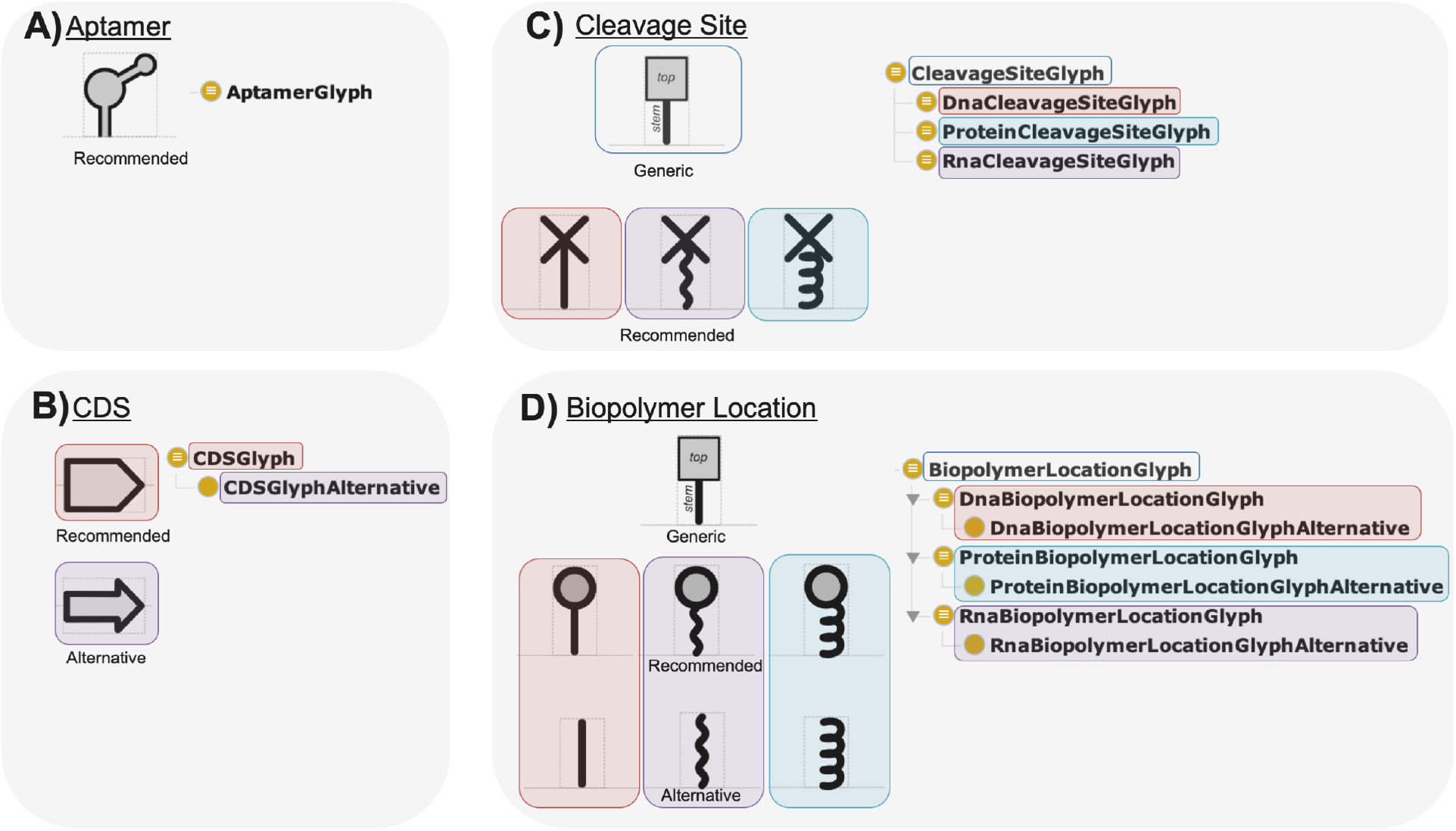
SBOL-VO captures relationships between glyphs that are defined as generic, recommended, or alternative. The ontology provides a unique term for each of these glyphs. Terms are decorated with ontological axioms to specify mapping rules for biological roles. Shown here are examples of A) a single glyph, B) a single-glyph and an alternative, C) multi-glyphs, D) multi-glyphs and alternatives

SBOL Visual also defines alternative glyphs. For example, although a recommended glyph is available for CDSs, users can also use an alternative glyph (Figure 2B). SBOL-VO provides terms for both recommended and alternative glyphs. Alternative terms are subclasses of recommended terms. Moreover, relationships between alternative and recommended terms are explicitly captured via *SomeValuesFrom* restrictions using the `isAlternativeOf` property. This property is not defined to be *functional*, i.e., a glyph term can be defined to be an alternative for multiple recommended terms. For example, regarding the ‘Complex’ category, SBOL-VO defines three recommended glyphs to represent a protein bound to a small molecule, a protein bound to a guide RNA, or a protein bound to another protein. For all three recommended terms, a single alternative term is defined. Recommended terms can also share a base term. For example, the base term for ‘Cleavage Site’ is used to describe generic shapes of multiple recommended cleavage sites for DNA, RNA, and protein molecules (Figure 2C). If alternatives are included for these recommended terms with generic parent terms, alternative terms are created as subterms (Figure 2D).

Ontological axioms restricting the use of glyphs for different sequence features, molecules, or molecular interactions can directly be utilised in ontological queries and be submitted to existing reasoners, such as HermiT.^11^ SBOL-VO incorporates these restrictions as defined classes for recommended terms, with alternative terms inheriting these restrictions. Glyphs representing genetic parts can be retrieved using the ‘`isGlyphOf some ComponentDefinition`’ ontological query. These queries can further be filtered with information about biological roles. For example, in order to retrieve glyphs representing promoter parts, the ‘`isGlyphOf some (ComponentDefinition and (role some SO:0000167))`’ query can be used. SBOL-VO is available as a Resource Description Framework (RDF) XML file and hence can also be queried using SPARQL graph patterns to retrieve more detailed information.^12^

We have made the information in SBOL-VO available via a web service, serving as a demonstration of how SBOL-VO can be used as a library simplifying the implementation of visual tools. The web service (see Availability) was developed as a RESTful application and provides various methods to access glyphs and query information. The web service either returns images directly or RDF graphs that are serialised using the JSON format. The three interactions supported by this service are:

- Serving glyphs: The web service acts as an image service and returns HTTP URLs for glyphs. For example, the `“/glyph/{ONTOLOGY_TERM}”` REST interface returns an image corresponding to an SBOL-VO, SO, or SBO term. If there is no exact match for an external term, then the closest parent term, for which a glyph is assigned, is substituted to find the most suitable recommended glyph. For example, the AptamerGlyph term (`/glyph/AptamerGlyph”`) from the SBOL-VO, or aptamer (`“/glyph/SO:0000031”`) or RNA_aptamer (`“/glyph/SO:0000033”`) terms from the SO can all be used to return the Aptamer glyph. PNG and SVG images can be explicitly retrieved by appending “/png” or “/svg” at the end of this interface, i.e. `“/glyph/SO:0000031/svg”`.
- Retrieving information about the mappings: The `“/mapping/{ONTOLOGY_TERM}”` REST interface is similar to the query interface but returns the mapping information rather than a glyph for a query term. For example both `“/mapping/SO:0000031”` and `“/mapping/SO:0000033”` returns information about the aptamer glyph. If a generic parent term is substituted, then the identifier of the parent term and its distance to the query term are also included in the results. The distance is measured as the number of child terms between the parent and query terms, including the parent term.
- Querying for glyphs: The `“/query/{ONTOLOGY_TERM}”` REST interface returns identifiers and information about how to locate glyphs linked to the provided SBOL-VO, SO, or SBO term and all its child terms. For example, both `“/query/CDSGlyph”` and `“/query/SO:0000316”` returns all SBOL-VO terms about the CDS glyph and its child terms, while `“/query/SBO:0000231”` returns information for all the glyphs linked to “molecular interactions”.
- Retrieving metadata: The `“/metadata/{SOL-VO_TERM}”` REST interface returns information about the provided SBOL-VO term. For example, `“/metadata/CDSGlyph”` returns information about the CDS glyph.

In summary, SBOL-VO makes standard glyphs used for genetic circuit diagrams available to computational tools in the form of an ontology. As a machine-accessible resource, SBOL-VO facilitates retrieving these glyphs unambiguously and allows queries to be submitted to existing reasoners. Since the ontology is available as an RDF graph, it can also be stored in triplestores or queried directly via SPARQL. This paper also presents a web service to resolve glyphs using HTTP, search for glyphs, and return detailed information about these glyphs, unifying access to standard glyphs via a REST-based HTTP interface. SBOL-VO thus facilitates integrating information, particularly for genetic design tools. The SBOL community heavily uses ontological terms to map genetic parts and their roles. Here, the creation of SBOL-VO and its mapping with the SBOL ontology facilitates further data integration for querying and retrieval of appropriate glyphs for genetic parts and their interactions. We note also that the construction of SBOL-VO is already automated. As a result, when future SBOL Visual specifications are released, new glyphs can easily be incorporated. SBOL-VO can act as a a catalogue that can be converted into different formats and hence can be used to autogenerate portions of the SBOL Visual specification in the future. Finally, further developments may also provide additional access methods to manipulate SBOL Visual glyphs in order to allow a greater degree of machine-assisted flexibility in visualization.

## Methods

The SBOL Visual 2 Ontology (SBOL-VO) was programmatically constructed using Markdown files that are created and managed by the SBOL Visual community. The programmatic conversion was carried out using the Python programming language and the Owlready API. ^13^ The HTML version of the ontology was generated using LODE. ^14^ Glyphs were then incorporated into the HTML documentation programmatically using Java. The SBOL-VO web service was developed as a RESTful^15^ application using Java and the Jersey (https://eclipseee4j.github.io/jersey) framework. The integration of ontologies and information retrieval were carried out using SPARQL and the Jena (https://jena.apache.org) framework.

## Availability

The SBOL Visual Ontology (SBOL-VO) and a human-browsable HTML catalogue of related terms and glyphs is available at http://synbiodex.github.io/SBOL-visual/Ontology.

The SBOL-VO web service is publicly available at http://vows.sbolstandard.org. Information about how to access different web service methods is also available from the same web page.

## Acknowledgement

J.B. is supported in part by NSF Expeditions in Computing Program Award #1522074 as part of the Living Computing Project. C.M. is supported by NSF grant #1939892. T.E.G. is supported by BrisSynBio, a BBSRC/EPSRC Synthetic Biology Research Centre grant BB/L01386X/1, and a Royal Society University Research Fellowship grant UF160357. G.-B.S. is supported by a UK EPSRC Fellowship for Growth in Synthetic Biology (EP/M002187/1) and a UK Royal Academy of Engineering Chair in Emerging Technologies for Engineering Biology. A.W. is supported by EPSRC grants EP/R003629/1 and EP/N031962/1. Any opinions, findings, and conclusions or recommendations expressed in this material are those of the author(s) and do not necessarily reflect the views of the funding agencies. This document does not contain technology or technical data controlled under either U.S. International Traffic in Arms Regulation or U.S. Export Administration Regulations.

## Notes

#### Summary of Updates

Updated the web service URL.

